# Automated Reconstruction of Whole-Embryo Cell Lineages by Learning from Sparse Annotations

**DOI:** 10.1101/2021.07.28.454016

**Authors:** Caroline Malin-Mayor, Peter Hirsch, Leo Guignard, Katie McDole, Yinan Wan, William C. Lemon, Philipp J. Keller, Stephan Preibisch, Jan Funke

## Abstract

We present a method for automated nucleus identification and tracking in time-lapse microscopy recordings of entire developing embryos. Our method combines deep learning and global optimization to enable complete lineage reconstruction from sparse point annotations, and uses parallelization to process multi-terabyte light-sheet recordings, which we demonstrate on three common model organisms: mouse, zebrafish, *Drosophila*. On the most difficult dataset (mouse), our method correctly reconstructs 75.8% of cell lineages spanning 1 hour, compared to 31.8% for the previous state of the art, thus enabling biologists to determine where and when cell fate decisions are made in developing embryos, tissues, and organs.

## 1 Main

With recent advances in light-sheet imaging techniques, it is possible to acquire whole embryo developmental datasets over long time scales with high spatial and temporal resolution in complex organisms such as mouse, *Drosophila*, and zebrafish (Wan et al., 2019a). The resulting datasets contain information required to track the movement and division of nuclei over time, yielding lineage trees and quantitative data on cellular dynamics that are crucial to the study of developmental biology at the cellular level (Spanjaard and Junker, 2017). However, manually tracing lineages with dedicated tools like MaMuT (Wolff et al., 2018) or Mastodon (https://github.com/mastodon-sc/mastodon) is arduous, and for complex, developing organisms it is only feasible to annotate a small percentage of all tracks, making automatic cell tracking necessary for holistic analysis.

Cell tracking algorithms have been developed for and tested on diverse datasets with different characteristics. While hand-engineered features are sufficient for cell detection and tracking in some model organisms (Bao et al., 2006; Amat et al., 2014), learned dataset-specific features, given sufficient training data, improve performance for datasets with heterogeneous cell or nucleus phenotypes and varying imaging statistics over time and space. In particular, deep learning has been shown to improve cell detection (Kok et al., 2020; Hayashida et al., 2020), segmentation (Weigert et al., 2020; Cao et al., 2020; Stringer et al., 2021; Medeiros et al., 2021), and tracking (Sugawara et al., 2021; Ulman et al., 2017; Moen et al., 2019; Hayashida et al., 2020; Medeiros et al., 2021) on a variety of datasets. Additionally, it has been shown that tracking methods that take into account global spatiotemporal context perform better, especially for datasets with more movement between time frames (Ulman et al., 2017). Tracking by graph optimization over a large spatiotemporal context allows inclusion of biological knowledge about track length and cell cycle, improving track continuity (Jug et al., 2016; Haubold et al., 2016; Kok et al., 2020) and even allowing recovery from noisy detection and segmentation (Schiegg et al., 2013).

Only a few of the aforementioned cell tracking methods are readily applicable to the unique challenges posed by contemporary 3D light-sheet datasets, the focus of this work. Practical methods for this kind of data should take into account temporal and 3D spatial context, easily scale to multi-terabyte datasets, and ideally should not require a manual segmentation of cells for training, due to the time required to generate per-pixel ground truth. Of methods that fulfill these requirements, Tracking with Gaussian Mixture Models (TGMM) (Amat et al., 2014) has been shown to work well on model organisms with approximately ellipsoid nuclei. More recently, the ELEPHANT tracking platform employed deep learning for cell detection and per-frame linking in light-sheet datasets with diverse cell appearance and movement (Sugawara et al., 2021). ELEPHANT requires a manual pseudo-segmentation of nuclei by ellipsoid fitting, which takes less time to generate than a per-pixel manual segmentation, but more than point annotations.

Our method combines global optimization and learned features, generating cell lineages through global graph optimization with learned costs. We show that this combination substantially decreases tracking error on three diverse datasets of different model organisms with different temporal resolution, signal to noise ratio, and nuclear appearance. Features are learned from sparse point annotations produced by current manual lineage tracking tools like MaMuT and Mastodon, and thus do not require a manual segmentation or dense lineage annotations, which allows rapid generation of training data. Crucially, the steps of our method—including the global optimization—can be computed in a distributed fashion, which is necessary to process multi-terabyte light-sheet datasets and enable the study of whole embryo morphogenesis.

An overview of our method is shown in Fig. 1. Because we are learning features from the data, the method is not tied to a specific input type or format: we use fused and unfused light-sheet recordings with a single fluorescent nuclear channel, and could easily extend to multi-channel input. We use sparse point annotations to train a convolutional neural network to predict at each pixel a *cell indicator* value that peaks at the center of each nucleus (Höfener et al., 2018; Kok et al., 2020), and a *movement vector* that points to the center of the same cell nucleus in the previous time frame (Hayashida et al., 2020; Sugawara et al., 2021). From these predictions, we generate a *candidate graph* in two steps: first, we place nodes at the local maxima of the cell indicator values to represent possible cell center locations, with a score to encode the network’s confidence. Second, we locally connect nodes in adjacent frames with edges to represent the possibility that the nodes represent the same cell, and assign a score to each edge based on agreement with the predicted movement vector.

**Figure 1:**
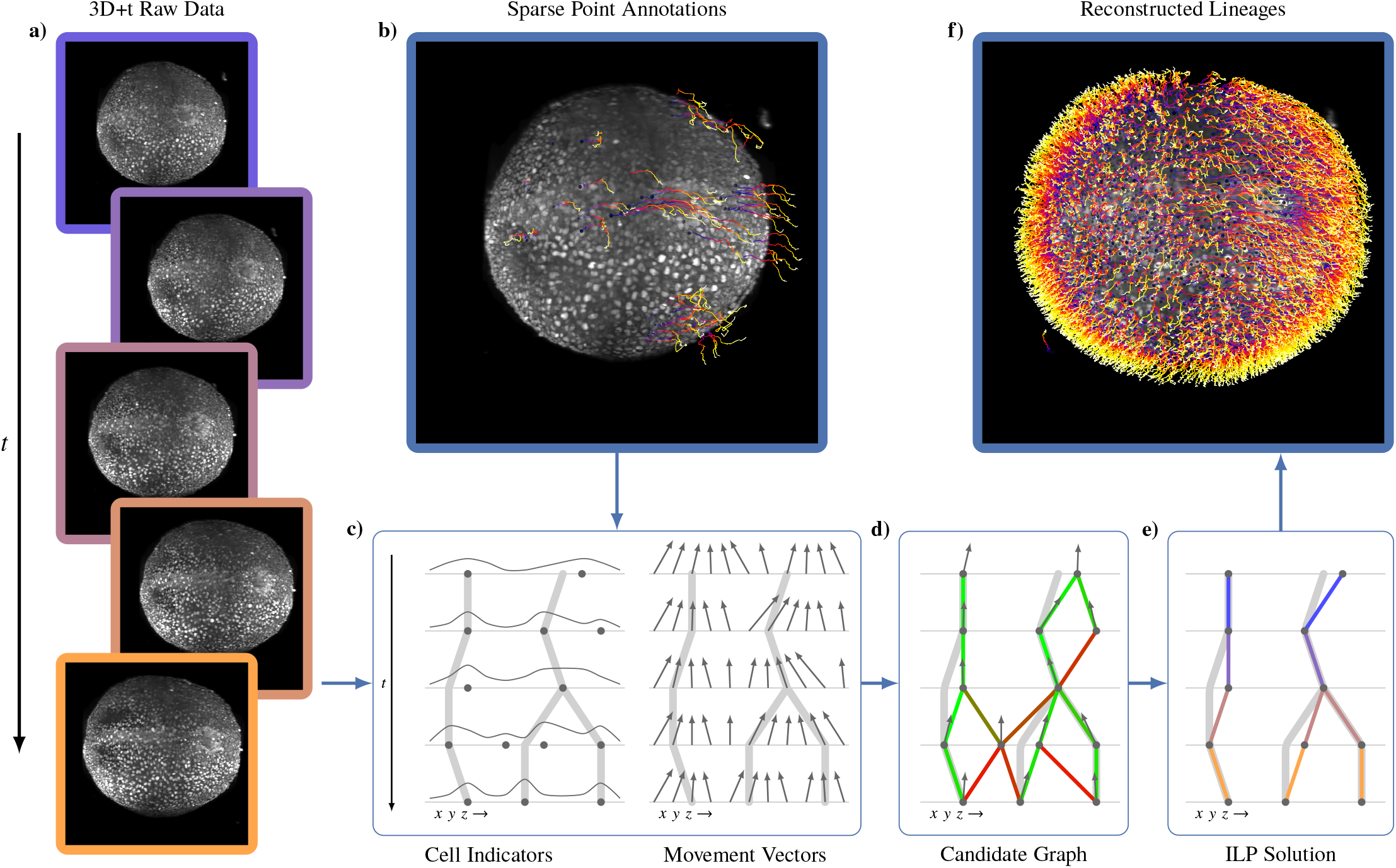
Overview of the method, including data and results from the mouse dataset. **a)** Raw mouse data over 50 time points, visualized as a max intensity projection. **b)** Sparse point annotations superimposed over the first frame of the raw data. Purple dots show the locations of annotated cells in the first time point, and the tails show the movement over time. **c)** Output of our cell indicator and movement vector networks. Light grey represents ground truth lineages. The cell indicator is trained to have maxima at the center of each nucleus, and the movement vector network is trained to predict the relative location of the same cell nucelus in the previous time point. **d)** Candidate graph extracted from the network output. Candidate cells are at cell indicator maxima, and nearby cells are connected with edges that are scored by agreement with the movement vector. **e)** Consistent lineage trees extracted from the candidate graph by global optimization using learned features and biological priors. **f)** Densely reconstructed lineages visualized over the mouse data.

Next, we solve a global constrained optimization problem on the candidate graph to select a subset of nodes and edges that form coherent lineage trees. We know that between time frames, cells can move, divide into two, enter or leave the field of view, or die, but not merge or split into more than two. Thus, we introduce hard constraints to prevent merging and divisions producing more than two progeny. The objective function incorporates prior knowledge that cell movement is much more common than division, death, and entering or leaving the field of view, encouraging long, continuous lineages by penalizing the start and end of tracks. These tree constraints and continuity costs are similar to those in previous work (Schiegg et al., 2013; Jug et al., 2016; Kok et al., 2020); however, we also incorporate the node and edge scores generated by the neural networks into the objective function as learned costs. Thus, we optimize for valid lineages that are both continuous and supported by the learned cell location and movement features. Our Integer Linear Program (ILP) formulation of the optimization problem additionally allows solving piece-wise in parallel on large datasets by introducing additional constraints to ensure consistent solutions between adjacent regions.

We evaluate our method on three sparsely annotated datasets from different commonly used model organisms to study embryogenesis: mouse (McDole et al., 2018), *Drosophila* (Amat et al., 2014), and zebrafish (Wan et al., 2019b) (see Supplementary Note 1 for details about the datasets and annotations). We compare the performance of our method against TGMM, the previous state-of-the-art method on these datasets (Amat et al., 2014; McDole et al., 2018), and greedy tracking using a per-frame nearest neighbor linking algorithm similar to the ELEPHANT tracking method (Sugawara et al., 2021). We compute multiple metrics, including the fraction of perfectly constructed lineages over a range of time periods, and errors per ground truth edge, broken into the following error types: false negative edges (FN), identity switches(IS)—when two tracks switch off following the same cell—false positive divisions (FP-D), and false negative divisions (FN-D), as illustrated in Fig. 3. False positive edges cannot be computed using sparse ground truth, because we cannot tell if unmatched reconstructions are false positives or tracking unannotated cells, and thus they are not included in our quantitative analysis. We show in Fig. 2 that, with around 20 hours of ground-truth annotation effort, our method correctly reconstructs more cell lineages than both baselines over all time ranges for all datasets. The largest improvement compared to TGMM is on the mouse dataset: our method correctly tracks 75.8% of mouse cells over a time span of 1 hour (12 time frames), compared to 31.8% for TGMM. By 175 minutes (35 frames), our method still correctly tracks more than half of all cells, while TGMM tracks less than 8%. On all three datasets, our method greatly reduces false negative edges compared to TGMM, while compared to the greedy baseline, our method produces far fewer false positive divisions. Supplementary Note 2 contains a detailed description of the evaluation metrics and baselines, and further observations about the performance on various metrics across organisms and evaluation regions.

**Figure 2:**
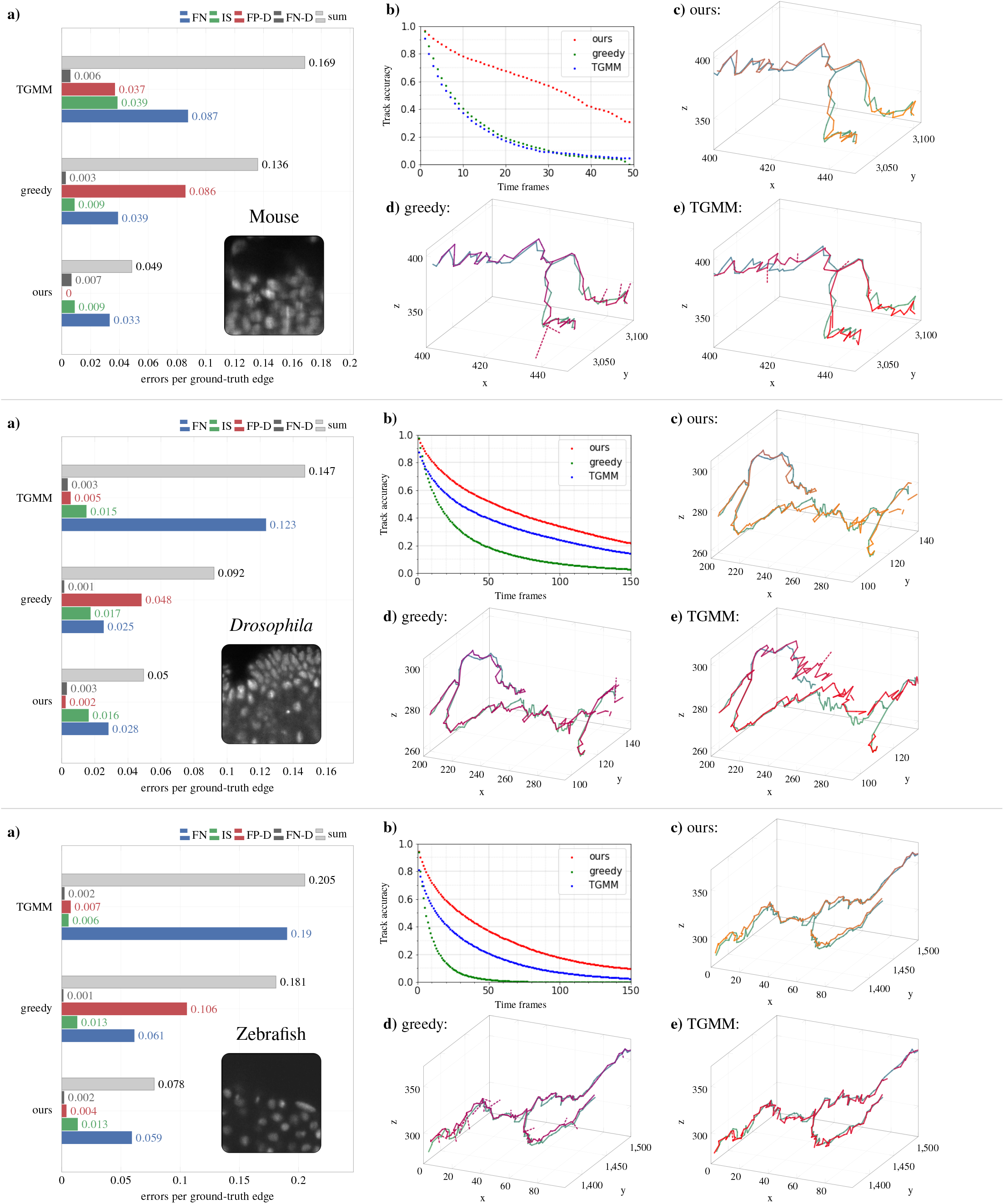
Comparison of tracking errors on three datasets (top to bottom: mouse, *Drosophila*, zebrafish). **a)** Average errors per ground-truth edge for each error type. **b)** Fraction of error-free tracks for a given track length. **c-e)** Example ground truth track (green) with superimposed tracking result (orange or red) for our method, the greedy baseline, and TGMM respectively. Other than the dashed false positive divisions, we only show detections that matched the selected ground truth track.

**Figure 3:**
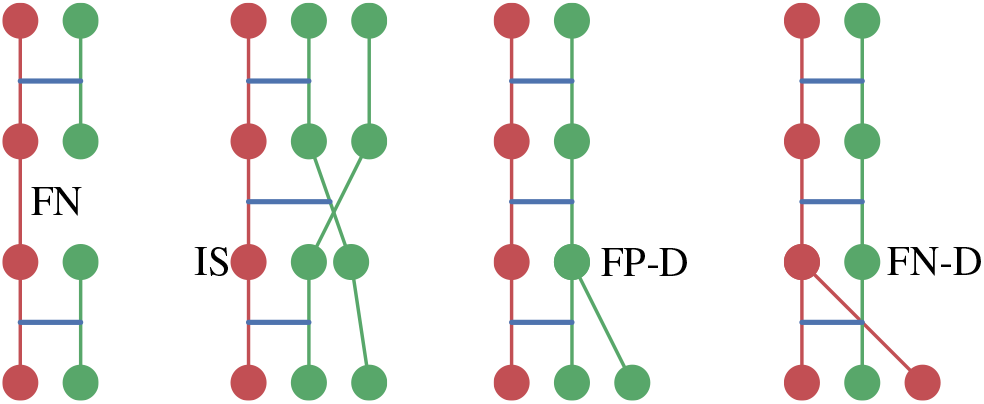
Diagram illustrating the four kinds of tracking errors used in our analysis: false negative edges (FN), identity switches (IS), false positive divisions (FP-D) and false negative divisions (FN-D). False positive edges are not pictured, as they cannot be determined from sparse ground truth. Red graphs represent ground truth tracks and green reconstructed tracks, while blue lines represent edges that are matched between the ground truth and reconstructed tracks.

Both the candidate graph generation and lineage optimization steps of our method are fully parallelizable and scale linearly with the size of the recording, which enables dense lineage reconstruction on very large datasets in reasonable time. On 20 GPUs and 100 CPU cores, reconstruction of dense lineages took about 44 hours on the 4.7TB mouse dataset, generating more than 7 million cell detections and 360,000 tracks over the 44 hour recording. Given the estimates that there are 6 million true cell detections in the dataset and that an annotator can click on a cell center every 1.5 to 3 seconds, it would take 2500-5000 annotator-hours to manually trace all lineages in this dataset. The source code of our method is publicly available, together with training and inference scripts and extensive documentation (https://github.com/funkelab/linajea).

The ability to densely reconstruct cell lineages in such large, information-rich datasets opens up vast opportunities for exploring cell fate dynamics and tissue morphogenesis. Accurately following cells and their progeny over extended time periods allows identification of individual cell behaviors that are not visible with shorter and less accurate lineages. For example, being able to accurately track more than half of all cells over a time window of 175 minutes in the mouse dataset, compared to only 30 minutes with previous methods, greatly reduces the manual curation needed to test hypotheses such as the existence of neuromesodermal progenitors which can produce neural or mesodermal progeny. While further work is required to improve cell division detection, the dense cell lineages we publish with this method are a rich source of information about the development and cell fate dynamics of common model organisms.

## 2 Method

### 2.1 Network Architecture, Training, and Prediction

To attain per-voxel predictions for cell locations and movements, we use a U-Net architecture with four resolution levels (Ronneberger et al., 2015). To incorporate temporal as well as spatial context, we concurrently feed seven 3D frames centered on the target time point and use four-dimensional convolutions until, due to valid convolutions, the time dimension is reduced to one. While we mostly downsample by (2,2,2), our mouse and *Drosophila* datasets are anisotropic, for the first pass we only downsample by 1 in z. We use 12 initial feature maps and increase by a factor of 3 at each level. When upsampling, we restrict our upsampling convolutional kernels to constant values, as we have observed this reduces artifacts in the output.

The cell indicator network is trained on sparse point annotations and predicts the centers of cell nuclei. The training signal for this network, called the cell indicator value, is a Gaussian with max value 1 at the cell center annotation and decreasing according to a hyperparameter σ. With only sparse annotations, it is unknown if pixels far from cell center annotations are background or cells that were not annotated. To avoid training on unknown regions, we construct a *training mask* around each annotation with a user-defined radius. This radius should be small enough that the mask will not overlap with neighboring cells. We only train on the mean squared error loss within the training mask. We are not training our cell indicator network on any background regions, so the behavior is unconstrained in the background. After prediction, we use local Non-Maximum Suppression (NMS) to extract cell center candidates, with the goal of detecting all cell centers along with potential false positives due to the unconstrained background behavior. The NMS window size is dataset dependent and should be a bit smaller than the minimal distance between two cell centers. To reduce the number of false positives, especially in background regions, we only consider detections with a minimal cell indicator value, determined empirically for each dataset and model. Additionally, if a foreground mask is available (as in the zebrafish dataset) we filter detections to those that lie in the foreground.

In addition to the cell indicator network, we train a movement vector network to predict the movement of cells between frames. For a pixel near to a cell in frame *t*, the movement vector is a 3D offset vector that points to the relative location of the center of the same cell in frame *t* − 1. Predicting the offset to the same cell in the previous time frame, rather than the next time frame, allows divisions to be represented naturally, since each daughter cell points to the center of the parent cell. We calculate the loss on two different masked regions. Loss *L*_*A*_ is the mean squared error between the ground truth and predicted movement vectors, calculated over the same training mask as the cell indicator network. Loss *L*_*B*_ limits the error to voxels with maximal cell indicator values after NMS that also are within the training mask. The total loss is the weighted sum *L* = *αL*_*A*_ + (1 − *α*)*L*_*B*_, with 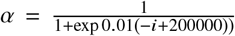. and *i* being the number of training iterations. This weighting scheme weights *L*_*A*_ higher at the beginning of training, when the cell indicator network is still converging, with a smooth transition to *L*_*B*_ at 200000 iterations.

These networks are trained simultaneously for 400000 iterations, with batch size 1. Batches are randomly sampled from annotated locations, and random augmentations including elastic deformation, mirroring, transposing axes, and intensity augmentation are applied using the Gunpowder library (https://github.com/funkey/gunpowder). Prediction is then performed blockwise using paralellization over multiple GPUs to process large datasets efficiently. To eliminate edge artifacts, we ensure that our prediction stride is a multiple of the network downsample factors (Rumberger et al., 2021).

### 2.2 Candidate Graph Extraction

After prediction, we create a directed candidate graph *G* = (*V, E)* with nodes that represent possible cell center locations and edges that represent possible movements of the same cell between frames. *G* is expected to contain extra nodes and edges, which will be filtered out in the final step.

*V* is the set of NMS detections. Each *v* ∈ *V* has a three dimensional location *l*_*v*_, a time *t*_*v*_, a predicted cell indicator score *s*_*v*_, and a predicted movement vector *m*_*v*_. We avoid storing the predicted cell indicators and movement vectors at every pixel by performing NMS on the cell indicator values during prediction and only saving the predicted values at the detection.

We construct the set of directed edges *E* by locally connecting nodes in adjacent frames with edges that point one frame backwards in time. For each candidate *v* at time *t*_*v*_, we compute the predicted location 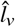 of the same cell in the previous frame: 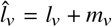. Then, we add an edge from *v* to each node candidate *u* at time *t*_*v*_ − 1 where the predicted distance 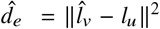 is less than a hyperparameter *θ*. 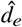 is stored as a score on each edge.

### 2.3 Discrete Optimization to Find Linage Trees

We construct a lineage tree by selecting a subset of nodes and edges from *G*. We define a vector ***y*** = [***y***^*V*^, ***y***^*E*^]^*T*^ ∈ {0.1}^|*V*|+|*E*|^ such that each element of the vector corresponds to a node or edge in *G*. Then *G*(**y**) is the subgraph induced by **y** that only contains nodes and edges with corresponding element of **y** equal to 1.

We then construct a constrained optimization problem that minimizes the objective

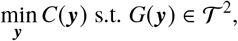

where 𝒯^2^ is the set of binary forests and *C* : **y** ⟶ ℝ assigns a cost for each set of selected nodes and edges. Thus, the goal is to select the cost-minimal subset of nodes and edges from *G* that form a binary forest.

To simplify the presentation of the cost function, we introduce two auxiliary indicator vectors of length |*V*| that can be entirely derived from **y**. The indicator for tracks appearing, **y** ^*A*^, is 1 for nodes at the beginning of a track and 0 otherwise. **y**^*D*^ represents a track disappearing and is 1 for leaf nodes at the end of a track. For a formalization of the definition of these auxiliary vectors from **y**, see Section 2.3.1.

With these auxiliary indicator variables, we define a linear cost function as follows:

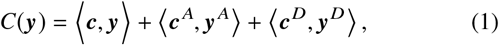

where **c** = [**c**^*V*^, **c**^*E*^ ]^*T*^ is a vector containing the cost for selecting each node and edge, and **c** ^*A*^ and **c**^*D*^ are vectors containing the cost of having a track appear (*c* ^*A*^) and disappear (*d* ^*A*^) at each node. The appear and disappear costs are constant hyperparameters of the method, but the predicted cell indicator values and movement vectors are used to individualize the cost vector **c** for selecting each node and edge.

With *s*_*i*_ as the cell indicator score for node *i*, we define the node selection cost for node *i* as 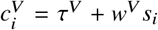, where *w*^*V*^ and τ^*V*^ are hyperparameters of the method. To encourage selection of higher cell indicator scores during minimization, *w*^*V*^ should be negative.

Similarly, with 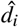 as the distance between the predicted and actual offsets at edge *i*, we define the edge selection cost for edge *i* as 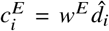. Unlike with node scores, *w*^*E*^ should be positive to encourage selection of edges with low scores, since those edges align better with the predicted cell movement.

To determine the optimal values of the ILP hyperparameters *c* ^*A*^, *c*^*D*^, τ^*V*^, *w*^*V*^, and *w*^*E*^, we performed a grid search where we fixed *c*^*D*^ = 1 to eliminate redundant solutions. We selected the hyperparameter set that minimized the sum of errors over the validation set (Supplementary Note 2).

#### 2.3.1 Integer Linear Program Formulation

We use an Integer Linear Program (ILP) to solve the constrained optimization problem with the Gurobi solver (Gurobi Optimizer, 2021). The objective is the cost function *C*(**y)** (Equation 1 in Section 2.3). To ensure a binary forest with correctly set auxiliary variables, we implement three kinds of constraints: consistency, continuation, and split constraints.

The consistency constraint requires that if an edge is selected, the incident nodes are selected as well. This constraint for edge *e* = (*v, u)* is represented by the equation 2*y*_*e*_ − *y*_*v*_ − *y*_*u*_ ≤ 0.

Two continuation constraints ensure that either the track continues or the node is marked as the beginning or end of the track. Let *P*_*v*_ be the set of edges from node *v* in time *t*_*v*_ to nodes in *t*_*v*_ − 1, and *N*_*v*_ be the set of edges from *t*_*v*_ + 1 to *v*. We define the appear continuation constraint as 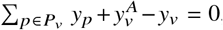 ensuring that if node *v* is selected, either there is one selected edge to time *t*_*v*_ − 1 or the appear indicator is set to 1. Additionally, we define the disappear continuation constraint as 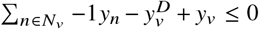 to ensure that either at least one edge to time *t*_*v*_ + 1 is selected or the disappear indicator is set to 1.

Finally, the split constraint ensures that the number of selected incoming edges is ≤ 2, *i.e*., for each node 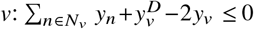.

#### 2.3.2 Processing Large Volumes Blockwise

Ideally, we would solve the ILP for the whole candidate graph at once to obtain a globally optimal solution. However, for large volumes this is too time and memory intensive. Therefore, to obtain lineage trees for arbitrarily large volumes, we divide the candidate graph into a set of blocks *B* that tile the whole volume and use multiple processes to solve the ILP for many blocks in parallel.

Solving each block *b* ∈ *B* completely independently can result in discontinuities in tracks between blocks, and the constraints would no longer be assured at the boundaries. To ensure a consistent, valid solution across the whole volume, we allow each process to view a context region around the target region *b* that must be at least as large as the amount a cell can move in space and time. Let 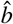 be the union of *b* and the surrounding context area. A process reads all nodes and edges in 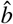, solves the ILP, and writes the result for only the target region *b* into a central database. If the database already contains results in the context region, these selections will be introduced as further constraints into the ILP, ensuring the solution will be consistent across boundaries. At the block boundaries, we set the appear and disappear costs to zero, because we do not want to penalize solutions that cross block boundaries.

The introduction of a context region introduces dependencies between neighboring blocks, and thus they cannot be run trivially in parallel. By ensuring that overlapping blocks are never run simultaneously using the Daisy library (https://github.com/funkelab/daisy), we ensure a valid, consistent global solution while retaining a high degree of parallel processing. While there is no guarantee of global optimality, with a large enough context region, we assume that nodes and edges further away do not affect the local solution in a target region.

## 3 Supplementary Note 1: Datasets and Annotations

We test our cell tracking method on time-lapse light-sheet recordings from three common model organisms: *Drosophila*, mouse, and zebrafish. We call these datasets Droso, Mouse, and ZFish, respectively, and summarize some relevant information about them in Table 1. Each dataset records a fluorescent nuclear marker: for ease of discussion, we will refer to each nucleus as corresponding to a single cell. While two of the datasets, Droso and Mouse, have a single view of the organism, the ZFish contains two orthogonal, registered but unfused views. To enable treating these views as interchangeable inputs to our networks, we resample them to isotropic resolution.

**Table 1:**
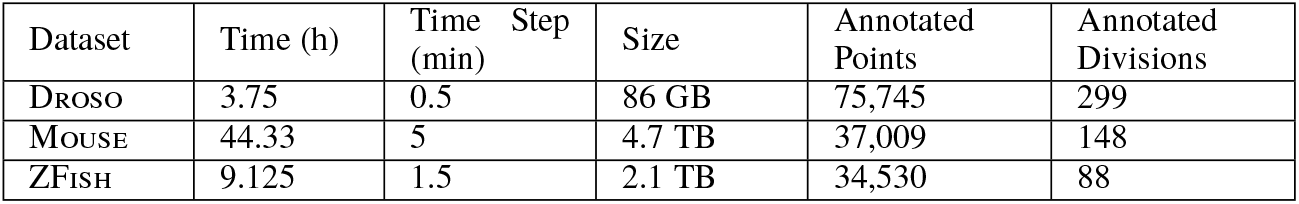
Summary information about the three datasets used to develop and evaluate our method.

We use sparse point annotations to train our method. As we leverage annotations originally performed for biological analysis, the annotated lineages are not randomly distributed, instead focusing on the developing nervous system of each organism. Although not necessary for training purposes, annotators ensured that lineages were fully traced by following a cell and all subsequent progeny until they were no longer visible. Thus, there are more annotations in later frames of each recording. See Table 1 for the number of cells and divisions annotated in each dataset.

For each organism, we divide the available annotations by time, location, and lineage into train, validation, and test sections, and report results on each split of the data individually, as well as averaged across splits as a form of k-fold cross validation. Cells in *Drosophila* and zebrafish rarely cross the center line of the organism, so we split the lineages into two groups based on side, discarding a small number of zebrafish lineages that did cross the center line. We then train two models using lineages from each side, leaving out 50 central time frames (200-250 for Droso, 150-200 for ZFish) for validation. We test each model on lineages from the side that was not used for training. Table 2 shows the number of cells and divisions in each train, validation, and test region for Droso and ZFish.

**Table 2:** Number of annotated cells (divisions) used for training, validation, and evaluation in Droso and ZFish. Sides of the organisms were arbitrarily labeled 1 and 2, and each split is named for the evaluation side.

| Split | Train | Validate | Test |
| --- | --- | --- | --- |
| DROSO side 1 | 30415 (96) | 7443 (54) | 38156 (149) |
| DROSO side 2 | 30839 (107) | 7596 (42) | 37589 (150) |
| ZFISH side 1 | 11919 (29) | 2159 (6) | 14535 (35) |
| ZFISH side 2 | 12281 (33) | 2346 (2) | 13996 (35) |

Within a developing mouse embryo, there is not a clearly defined center line that cells do not cross. Thus, instead of splitting lineages into groups by region, we define three sections of Mouse by time frame: “early” (50-100), “middle” (225-275), and “late” (400-450). Due to extensive embryonic development over the 44 hour recording, early, middle, and late stages represent different cell environments and organization, and there are far more cells by the end of the recording than at the early stages. Each model is trained leaving out two of those sections, one for validation and one for testing. This is repeated for all combinations of validation and testing, resulting in six total train/validation/test splits. The number of cells and division in each Mouse split is shown in Table 3.

**Table 3:** Number of annotated cells (divisions) used for training, validation, and evaluation in Mouse. Splits are named for the evaluation set, so early 1 and early 2 are both evaluated on the early section.

| Split | Train | Validate | Test |
| --- | --- | --- | --- |
| early 1 | 33744 (132) | 3178 (13) | 309 (3) |
| early 2 | 30509 (109) | 6413 (36) | 309 (3) |
| middle 1 | 33744 (132) | 309 (3) | 3178 (13) |
| middle 2 | 27640 (99) | 6413 (36) | 3178 (13) |
| late 1 | 30509 (109) | 309 (3) | 6413 (36) |
| late 2 | 27640 (99) | 3178 (13) | 6413 (36) |

## 4 Supplementary Note 2: EValuation

### 4.1 Metrics

Cell lineages can be used for a wide variety of analyses, and different kinds of errors can affect downstream results differently; therefore, reducing performance of a tracking method to a single number that represents “overall performance” is generally not possible. Therefore, we distinguish five types of tracking errors: false positive edges (FP), false negative edges (FN), identity switches (IS)—when one reconstructed track takes over following a cell from another reconstructed track—false positive divisions (FP-D) and false negative divisions (FN-D), as shown in Fig. 3. To allow comparison across datasets, we normalize the number of errors by the number of ground truth edges, resulting in an *errors per edge* metric. Additionally, we compute the fraction of ground truth lineages that were perfectly reconstructed over T time points, for a range of values for T. Ground truth segments over time T were identified using a sliding window of time T over the whole evaluation region, splitting at divisions only when they occur in the first frame of the window. Matched reconstructions were considered perfect when none of the error types above occurred over the course of the window.

Evaluating with sparse point annotations presents two unique challenges. First, we cannot determine false positive edges, so we omit this error type from our analysis. Due to the non-maximal suppression window used when extracting cell candidates, our method cannot naively minimize the false negative edge metric by extreme overdetection of false positive cells. However, false positive tracks can still appear, and without dense annotations we are limited to qualitative analysis. Second, we cannot use segmentation overlap to match ground truth to reconstructed cells. Instead, we choose a matching threshold that is a bit larger than the radius of a nucleus in the dataset. Considering only nodes within this threshold as potential matching endpoints, we pair ground truth and reconstruction edges using Hungarian Matching to minimize the sum of endpoint distance. Both reconstruction and ground truth edges can be matched to a dummy edge, allowing detection of false negative ground truth edges and reconstructions that do not match to any ground truth.

In addition to evaluating tracking performance, we examine the performance of the cell indicator and movement vector networks. The efficacy of the cell indicator model is determined by the cell recall, or the percent of ground truth cells that have a cell indicator maxima within the matching threshold. To evaluate the quality of the movement vectors, we find the closest cell indicator maxima for each ground truth node (within the matching threshold) and compute the distance between the parent location predicted by the movement vector at that maxima and the actual parent location. We use the “no movement” prediction as a baseline, to simulate the assumption that cells stay in the same place.

### 4.2 Baselines

Due to the size of our datasets and nature of our ground truth, we can only compare against cell tracking methods that can be run efficiently on multi-terabyte 3D datasets, and that do not require dense annotations or segmentations for training. Tracking with Gaussian Mixture Models (TGMM) (Amat et al., 2014) was previously run on certain time regions of Droso and Mouse, and we were able to extend those results to the full time series. Because TGMM cannot process multi-channel input, for ZFish we produced tracks for each of the two views separately, and reported the best result for each evaluation region. More recently, the tracking method included in the ELEPHANT framework has potential to be scalable to multi-terabyte datsets (Sugawara et al., 2021).

The cell detection step requires sparse nuclear segmentations by manual ellipsoid fitting, preventing us from comparing directly with the full method, so instead we run a greedy nearest-neighbor linking algorithm inspired by this work on our cell candidates. Starting in the final frame *t*, we consider all cell candidates to be part of a track. We then greedily select edges from *t* to *t* − 1 with the smallest difference between predicted and actual offset, enforcing the constraint that cells cannot divide into more than two by removing edges that connect to nodes in *t* − 1 that already have two selected incoming edges. We then process each subsequent pair of frames going back in time, first extending existing tracks, and then creating new tracks if any valid edges remain.

### 4.3 Results

Fig. 2 shows the sum of errors per edge for each organism, averaged over the train/validation/test splits. Across all datasets, our method produces significantly fewer errors per edge than TGMM and the greedy baseline, with the greedy baseline landing between TGMM and our method. Our method performs similarly between Mouse and Droso (0.049 and 0.050 total errors per edge) and slightly worse on ZFish (0.078).

Considering individual types of errors provides more insight into the performance of the different methods. For both TGMM and our method, false negative edges (FN) are the most common error type. In every case, our method produces fewer FN and fewer false positive divisions (FP-D) than TGMM. The false negative division (FN-D) performance is similar between our method and TGMM - in absolute numbers, neither our method nor TGMM correctly identifies more than a third of the divisions, but divisions are so underrepresented in the evaluation sets that this error type does not significantly affect the overall sum of errors. On Mouse, our method does not produce any divisions, and thus the FP-D rate is always zero, while TGMM has a very high FP-D rate and still does not detect many of the true divisions.

While our method does not always have fewer identity switches (IS) than TGMM, exmaning performance by dataset shows clear trends. For Mouse, our method always produces fewer IS than TGMM. However, for Droso and ZFish, TGMM produces hardly any IS, likely due to the high number of FN. Since an IS can only occur when two neighboring ground truth edges are matched to different reconstructed tracks, a high number of FN reduces the opportunities for IS to occur. Our method significantly reduces the number of FN on these datasets, resulting in slightly more IS than TGMM but fewer overall errors.

Fig. 2 also shows the track accuracy, or fraction of perfectly reconstructed tracks, for a range of track lengths. For length 1, this metric is the fraction of false negative edges, and thus our method and greedy outperform TGMM. However, as the time window increases, the greedy method’s track accuracy drops quickly for all datasets, reflecting the lack of global track optimization in this per-frame tracking algorithm. TGMM’s accuracy is similar to the greedy baseline on Mouse, but for Droso and ZFish, the slope of the decline is much flatter, indicating that TGMM perfectly reconstructs more long track segments on these datasets. Combining the fairly high track accuracy of TGMM with the large number of false negative edges, we can infer that on Droso and ZFish, TGMM’s errors are grouped together, resulting in some tracks being faithfully reconstructed and others missed completely. Our method has the highest track accuracy across all datasets, with a similar rate of decline as TGMM on Droso and ZFish but a higher starting accuracy.

To examine differences in performance between models trained and evaluated on tracks from different regions, we show results for each train/test split described in Supplementary Note 1 in Fig. 4 (errors per edge) and Fig. 5 (track accuracy). Due to the cross validation used for Mouse, we have results from two models for each evaluation region, with each model trained and validated on different data splits. For both sum of errors and track accuracy, the models that were trained on more data (early 1, middle 1, and late 1 as shown in Table 3) performed slightly better than those trained on less. The sum of errors also slightly increased for our method from early to late regions on Mouse, reflecting increasing difficulty of the task over time, although overall trends about relative performance and error types between our method and the baselines hold. Droso shows similar results between the two evaluation regions, but the same is not true for ZFish. TGMM performs much better on ZFish side 2 than side 1 in both track accuracy and sum of errors. Indeed, on side 2, TGMM and our method have a similar track accuracy, while TGMM performance on side 1 degrades significantly using both metrics. Manual examination of the raw data shows that the tracks on side 1 are harder for a human to identify due to less clear signal on that side of the dataset; thus, the relative results indicate that our method is more robust to varied imaging conditions than TGMM. Unexpectedly, the greedy baseline performs worse on the easier side 2. To explain this, we observe that the cell indicator model for side 2 predicts significantly more candidate cells, and more false positive candidates, than the model for side 1, likely due to randomness in the training pipeline (see Supplementary Note 3). The greedy method creates many false positive divisions involving those candidates, while ours does not, showing that the optimization step can extract coherent tracks from a noisy candidate graph.

**Figure 4:**
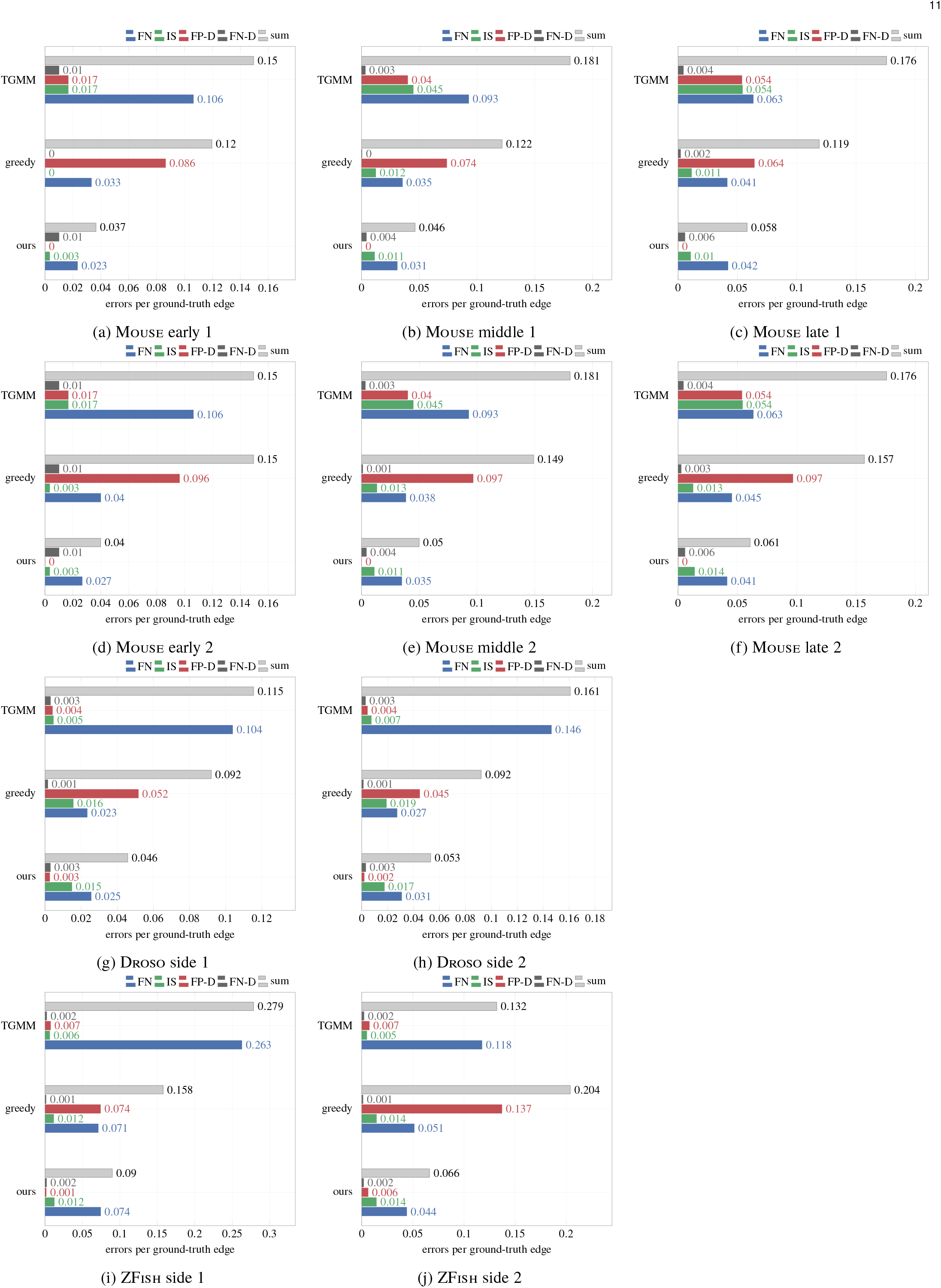
Errors per edge for our method, the greedy baseline, and TGMM, on all datasets and evaluation regions.

**Figure 5:**
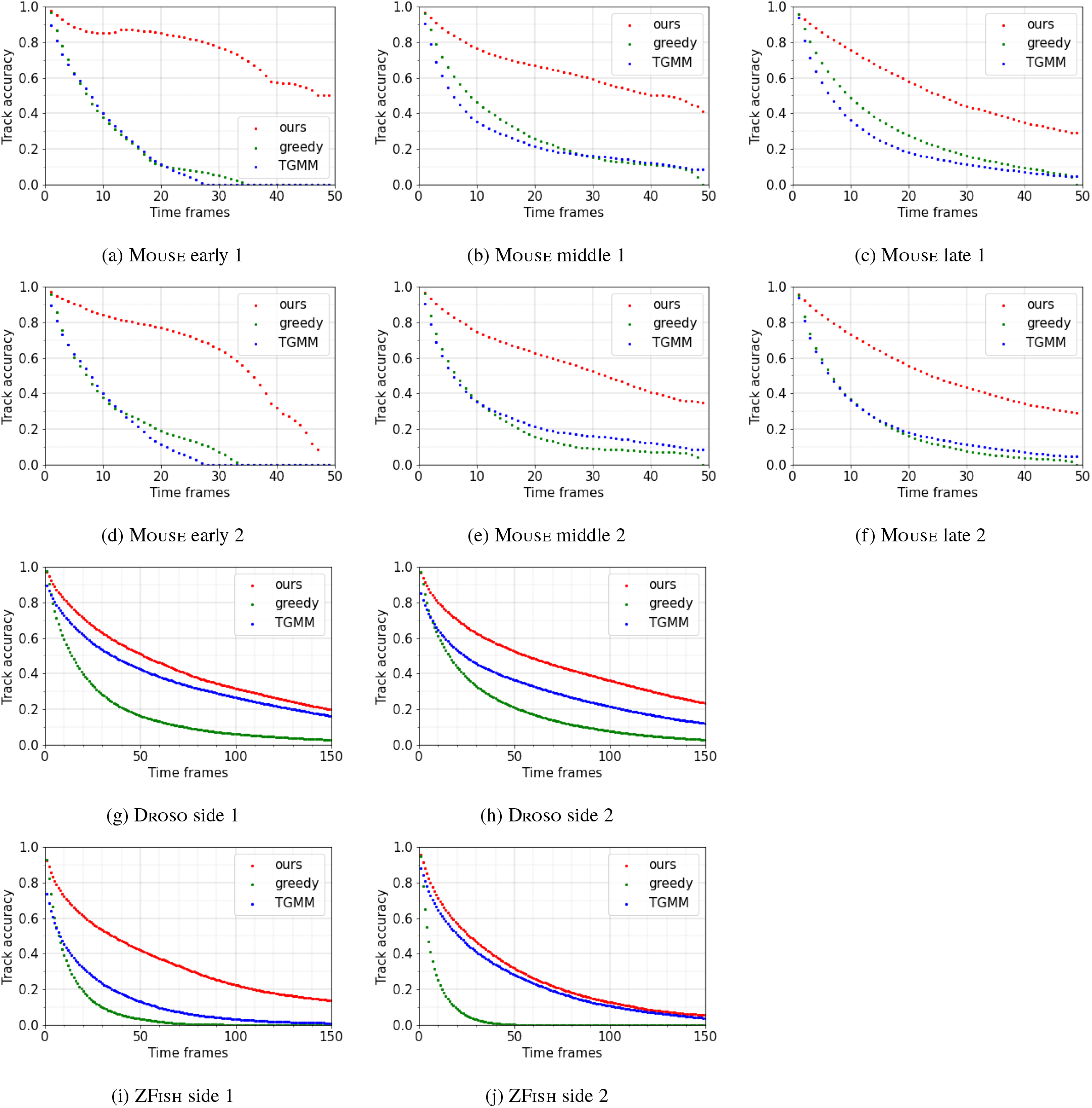
Fraction of ground truth segments correctly reconstructed over t time frames, for a range of t, on all datasets and evaluation regions.

Fig. 6 shows the standalone performance of the cell indicator and movement vector networks. Cell recall for all mouse and *Drosophila* models exceeds 0.99, indicating that nearly all ground truth cells in these datasets have a nearby candidate cell. Recall is slightly lower for both zebrafish models, but still exceeds 0.96. The movement vector network has a smaller mean distance between predicted and actual parent location than the baseline for all models, and the distribution of distances is concentrated closer to zero. The Mouse cells move further on average than the *Drosophila* and zebrafish cells, so the magnitude of improvement compared for Mouse is greater. The max distance is higher for our model than the baseline in all but one case, indicating that in the scenarios where cells move the most, such as after division, the movement vector network can point the wrong way. However, overall the movement vector network tends to point in the direction of the parent, as expected.

**Figure 6:**
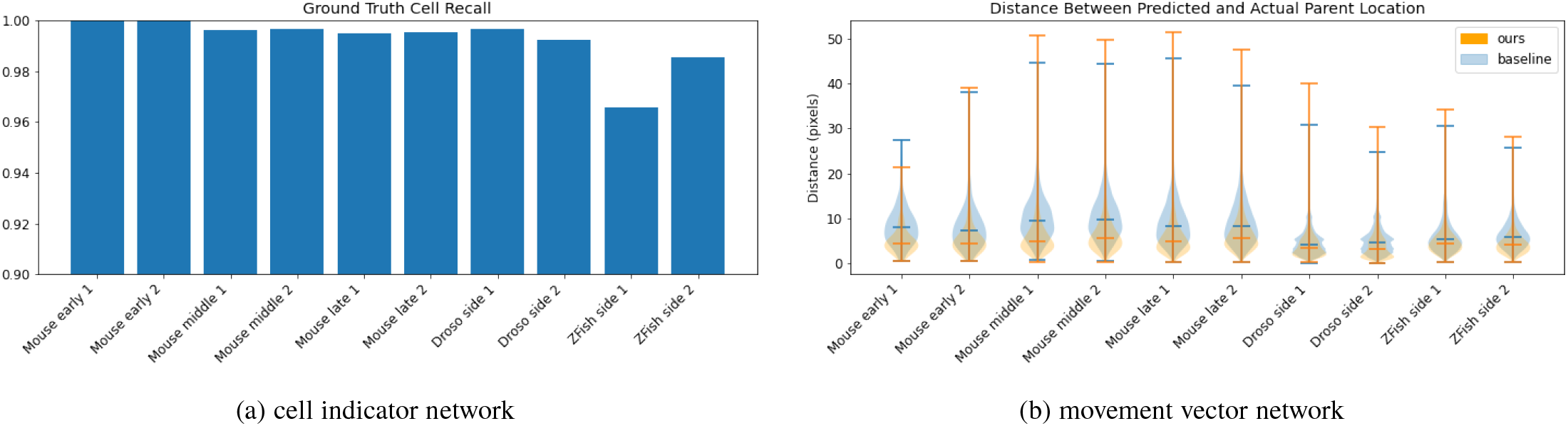
Performance of the cell indicator and movement vector networks. **(a)** Recall of cell indicator networks, as measured by the number of ground truth annotations in the evaluation set that have a cell indicator maxima within the matching threshold. **(b)** The distance between the predicted parent locations and actual parent locations for each ground truth cell with a matched candidate within the matching threshold, represented as a violin plot with hashes at the min, max and median values. Baseline of no movement is shown in blue, and our predicted movement vectors are shown in orange.

## 5 Supplementary Note 3: Ablation Study

Our training method contains multiple sources of randomness, from batch sampling to augmentation. To determine the effect of this randomness, we train, validate, and test the same model five times. The results shown in Fig. 7a illustrate that random batch selection and augmentation in training do affect tracking performance. The sum of errors and distribution of errors between false negative edges and identity switches vary substantially between the five models.

**Figure 7:**
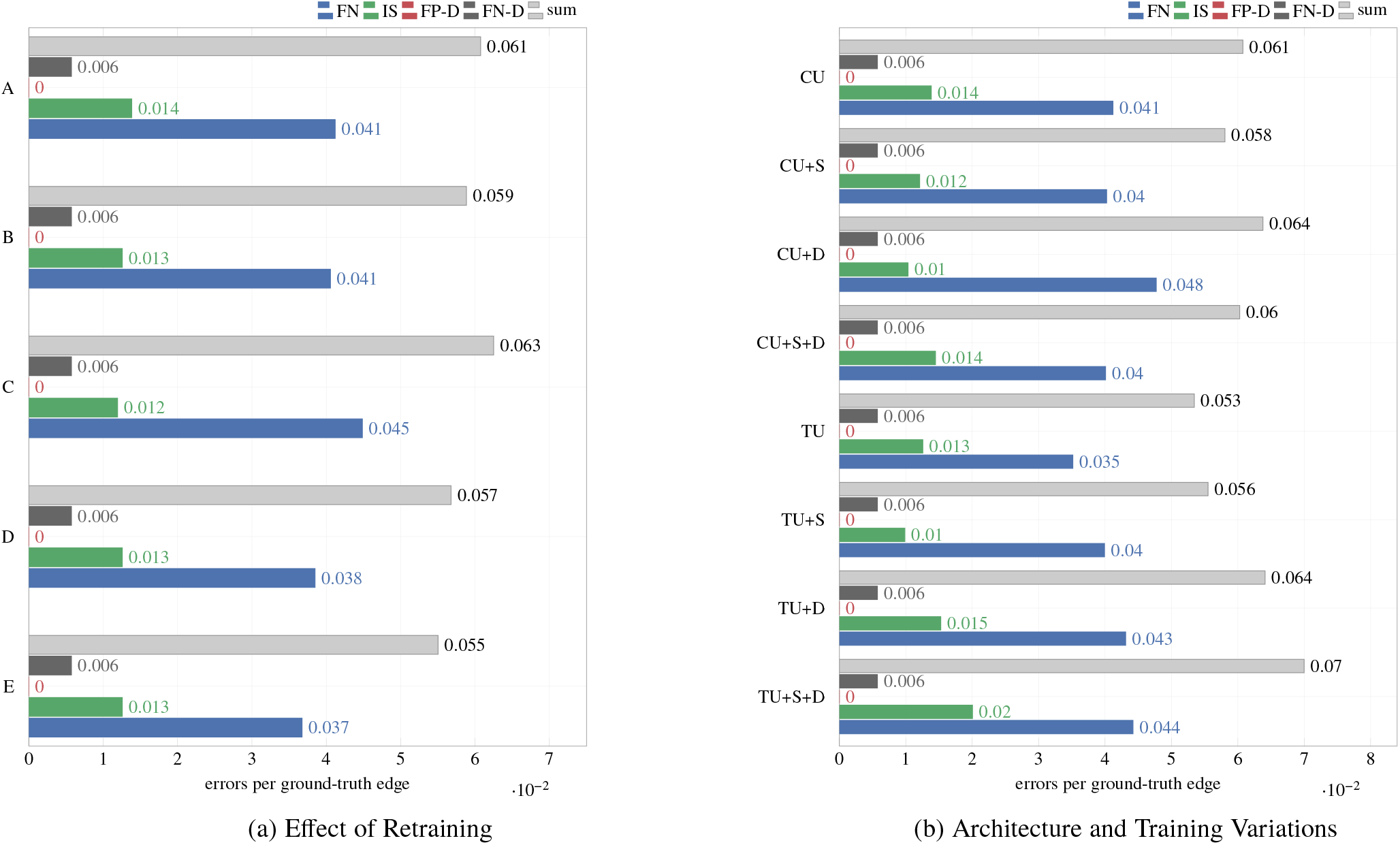
Supplementary experiments to determine the effect of randomness in training and architecture and training variations. **(a)** Sum of errors per ground truth edge, for five copies of the same model trained, validated, and tested on Mouse late 2. Variation in the error counts stems from random network initialization and random training sample selection and augmentation. **(b)** Sum of errors per ground truth edge for eight different models trained, validated, and tested on Mouse late 2, comparing constant (CU) and transpose upsampling (TU), shift augmentation (+S), and division sampling (+D).

In addition to our standard model, we test three changes to architecture, sampling, and augmentation. In our U-Net architecture, we experiment with two different upsampling methods: with and without limiting the transpose convolutional kernel to a kernel of ones. We call these two upsampling methods transpose upsampling (TU) and constant upsampling (CU). Because divisions are underrepresented in the training data and particularly difficult, we try sampling batches specifically at divisions 25% of the time (+D). We also simulate more cell movement by adding a random shift augmentation between the previous frame and the target frame (+S).

Results for each combination of these training and architecture variations on one Mouse split are shown in Fig. 7b. To draw conclusions about the effect of any of these variations, the resulting change in performance has to be greater than the effect of random retraining shown in Fig. 7a. In general, none of the models produced a large, consistent difference in tracking score, although division sampling seems to produce worse results in general. Due to training time and expense, we were not able to train every model in the ablation study multiple times or on every dataset, which would have allowed more conclusive comparisons. While further exploration into architecture and training decisions could yield incremental improvements, these initial results are insufficient to incorporate any of the three changes into our main model.

## 6 Supplementary Discussion

Using deep learning allows the method to adapt to different imaging conditions and organisms, and boosts performance compared to a heuristic approach as shown by the comparative performance of TGMM and the greedy baseline. However, deep learning in general requires annotated training data, which can be time consuming and costly to acquire. Our method minimizes the annotation burden by leveraging sparse point annotations in segments as short as two frames. Furthermore, the amount of training data required is reduced because the models do not need to achieve perfect performance: the global optimization can filter out superfluous detections and ignore individual inaccuracies in favor of global evidence and biological priors. Based on our results, between 10 and 30 thousand sparse point annotations created with MaMuT or Masodon would be sufficient to train a model to track cells in new organisms or imaging conditions. Assuming each point annotation can be generated in 3 seconds, sufficient training data can be produced in 8 to 24 hours of manual annotation.

While the optimization step filters out some false positive candidate detections in the backround, even better performance could be achieved by including negative examples in training. While we cannot quantitatively measure false positives when evaluating with sparse annotations, qualitative analysis of the results showed that the cell indicator network does predict false positives, especially in regions with high intensity but no discernible nuclei. When these false positive candidates persist through multiple frames, they can be linked to create false positive tracks. Incorporating our cell indicator model into the ELEPHANT interactive training and annotation framework (Sugawara et al., 2021) is a possible solution, since annotators could easily generate negative training examples in regions where the network most needs guidance. Using targeted negative examples, we expect the cell indicator network could learn to suppress cell prediction in these high intensity regions with limited fine-tuning.

When applying our method to a new dataset, one bottleneck of the method is the need to grid search the hyperparameters of the ILP. Even with blockwise processing, solving the ILP on the whole validation set takes tens of minutes per run, so we limited the grid search to four values per hyperparameter, resulting in 256 runs. We were guided by experience in choosing the range of values to search, but there is no guarantee that our solution was optimal, or that the same range would apply to different datasets. In the future, we will examine alternatives to grid searching a manually selected range of values, such as using a structured support vector machine to find the best set of ILP parameters on a given dataset.

Finally, identifying divisions is an important question for developmental analysis, but divisions are underrepresented compared to non-dividing cells, and have distinct movement and appearance. To address the difficulties that divisions present, we tried sampling divisions more frequently during training and adding a shift augmentation to mimic the movement of dividing cells, as discussed in Supplementary Note 3. However, even with these tactics, our method does not consistently identify divisions. One possible explanation is that the failure to identify divisions occurs in the optimization step, while our interventions focus on improving network predictions. During validation, we choose the ILP hyperparameters that minimize the sum of all error types. The underrepresentation of divisions in the validation set means that false negative divisions do not contribute heavily to the sum compared to false negative edges or even false positive divisions. Minimizing sum of errors thus can lead to models that select very few divisions, as long as the other error categories are minimized. Focused efforts to improve division performance will be necessary to attain reliable results.

## Acknowledgements

We thank William Patton and Tri Nguyen for supporting the Gun-powder and Daisy libraries, Steffen Wolf for his guidance and feedback, and Nils Eckstein, Julia Buhmann, and Arlo Sheridan for helpful discussions. **Funding** This work was supported by Howard Hughes Medical Institute. K.D. was supported by the Medical Research Council, as part of United Kingdom Research and Innovation [MCUP1201/23]. P.H. was supported by HFSP grant RGP0021/2018-102, the MDC Berlin-New York University exchange program, and the HHMI Janelia Visiting Scientist Program. **Author contributions** *Conceptualization*: Jan Funke, Philipp J. Keller. *Funding acquisition*: Jan Funke, Philipp J. Keller, Stephan Preibisch, Katie McDole, Peter Hirsch. *Software*: Caroline Malin-Mayor, Peter Hirsch, Jan Funke, Leo Guignard. *Validation and evaluation*: Caroline Malin-Mayor *Data and annotation generation*: Katie McDole, Yinan Wan, William C. Lemon. *Supervision*: Jan Funke, Stephan Preibisch, Philipp J. Keller. *Writing - original draft*: Caroline Malin-Mayor, Jan Funke. *Writing - review & editing*: Caroline Malin-Mayor, Jan Funke, Philipp J. Keller, Katie McDole, Stephan Preibisch, Peter Hirsch, Leo Guignard.

